## Supplements for "Improved gene regulatory network inference from single cell data with dropout augmentation"

### Supplements for Dropout Augmentation and DAZZLE

**Table S1. Performance on additional benchmarks**

| | hESC | hHep | mDC | mESC | mHSC-E | mHSC-GM | mHSC-L |
|---|---|---|---|---|---|---|---|
| # of Genes | 1410 | 1448 | 1321 | 1620 | 1204 | 1132 | 692 |
| # of Cells | 758 | 425 | 383 | 421 | 1071 | 889 | 847 |
| **Metric: Early Precision Ratio; Ground Truth: STRING** | | | | | | | |
| # of True Edges | 5,149 | 9,000 | 5,898 | 8,479 | 1,826 | 1,311 | 154 |
| GENIE3 | 3.66 (0.07) | 3.17 (0.03) | 2.19 (0.04) | 3.48 (0.03) | 7.00 (0.06) | 8.53 (0.09) | 7.34 (0.14) |
| GRNBoost2 | 3.46 (0.09) | 2.78 (0.04) | 2.08 (0.04) | 3.69 (0.05) | 6.21 (0.06) | 7.32 (0.12) | 7.12 (0.24) |
| DeepSEM-1x | 4.23 (0.12) | 3.12 (0.18) | 2.06 (0.09) | 3.89 (0.10) | 7.36 (0.22) | 7.91 (0.33) | 7.21 (0.47) |
| DeepSEM-10x | 4.54 (0.11) | *3.58 (0.18)* | *2.20 (0.07)* | *4.15 (0.07)* | 7.86 (0.07) | *8.96 (0.19)* | *7.43 (0.20)* |
| DAZZLE-1x | 5.12 (0.14) | 3.25 (0.08) | 2.02 (0.05) | 3.90 (0.08) | 7.79 (0.17) | 8.54 (0.15) | 7.37 (0.13) |
| DAZZLE-10x | *5.25 (0.05)* | 3.33 (0.03) | 2.01 (0.03) | 3.96 (0.07) | *8.06 (0.06)* | 8.53 (0.08) | 7.40 (0.07) |
| **Metric: Early Precision Ratio; Ground Truth: Non-celltype-specific ChIP-Seq** | | | | | | | |
| # of True Edges | 4,597 | 5,335 | 3,918 | 8,030 | 1,960 | 1,358 | 317 |
| GENIE3 | 1.68 (0.03) | 1.85 (0.08) | 2.93 (0.08) | 3.47 (0.05) | 4.64 (0.07) | 5.86 (0.09) | 3.05 (0.09) |
| GRNBoost2 | 1.49 (0.05) | 1.70 (0.05) | 2.58 (0.07) | *3.51 (0.05)* | 4.71 (0.11) | 5.03 (0.15) | 3.02 (0.11) |
| DeepSEM-1x | 2.20 (0.19) | 2.56 (0.12) | 3.29 (0.14) | 3.31 (0.12) | 5.55 (0.39) | 5.66 (0.32) | 3.14 (0.28) |
| DeepSEM-10x | 2.18 (0.05) | 2.67 (0.07) | *3.73 (0.08)* | 3.50 (0.06) | 6.04 (0.14) | 6.13 (0.22) | *3.35 (0.16)* |
| DAZZLE-1x | 2.31 (0.12) | 2.77 (0.16) | 3.51 (0.11) | 3.21 (0.14) | 6.29 (0.13) | 6.11 (0.11) | 3.30 (0.17) |
| DAZZLE-10x | *2.40 (0.04)* | *2.83 (0.05)* | 3.64 (0.07) | 3.26 (0.04) | *6.36 (0.06)* | *6.21 (0.05)* | 3.34 (0.06) |
| **Metric: Early Precision Ratio; Ground Truth: Celltype-specific ChIP-Seq** | | | | | | | |
| # of True Edges | 7,050 | 15,410 | 1,193 | 42,795 | 21,975 | 14,135 | 5,180 |
| GENIE3 | 0.95 (0.01) | *1.12 (0.01)* | 0.99 (0.06) | *1.06 (0.00)* | *1.02 (0.00)* | 1.03 (0.09) | *1.07 (0.00)* |
| GRNBoost2 | 0.78 (0.02) | 0.27 (0.00) | 0.97 (0.08) | 0.33 (0.00) | 0.21 (0.00) | 0.23 (0.01) | 0.44 (0.01) |
| DeepSEM-1x | 1.13 (0.04) | 1.03 (0.01) | 1.11 (0.07) | 1.02 (0.01) | 1.01 (0.00) | 1.02 (0.00) | 1.04 (0.01) |
| DeepSEM-10x | 1.15 (0.02) | 1.04 (0.00) | 1.15 (0.03) | 1.02 (0.01) | 1.01 (0.00) | *1.03 (0.00)* | 1.04 (0.00) |
| DAZZLE-1x | 1.20 (0.03) | 1.03 (0.01) | 1.11 (0.06) | 1.02 (0.01) | 1.01 (0.00) | 1.02 (0.00) | 1.06 (0.01) |
| DAZZLE-10x | *1.20 (0.01)* | 1.03 (0.00) | *1.17 (0.02)* | 1.02 (0.00) | 1.01 (0.00) | 1.03 (0.00) | 1.06 (0.00) |

Number of target genes: 1000.
Higher ratios indicate better performance. Here, italicized cell in dark shades indicate the best algorithms and lightly shaded cell indicate the 2nd best algorithm.

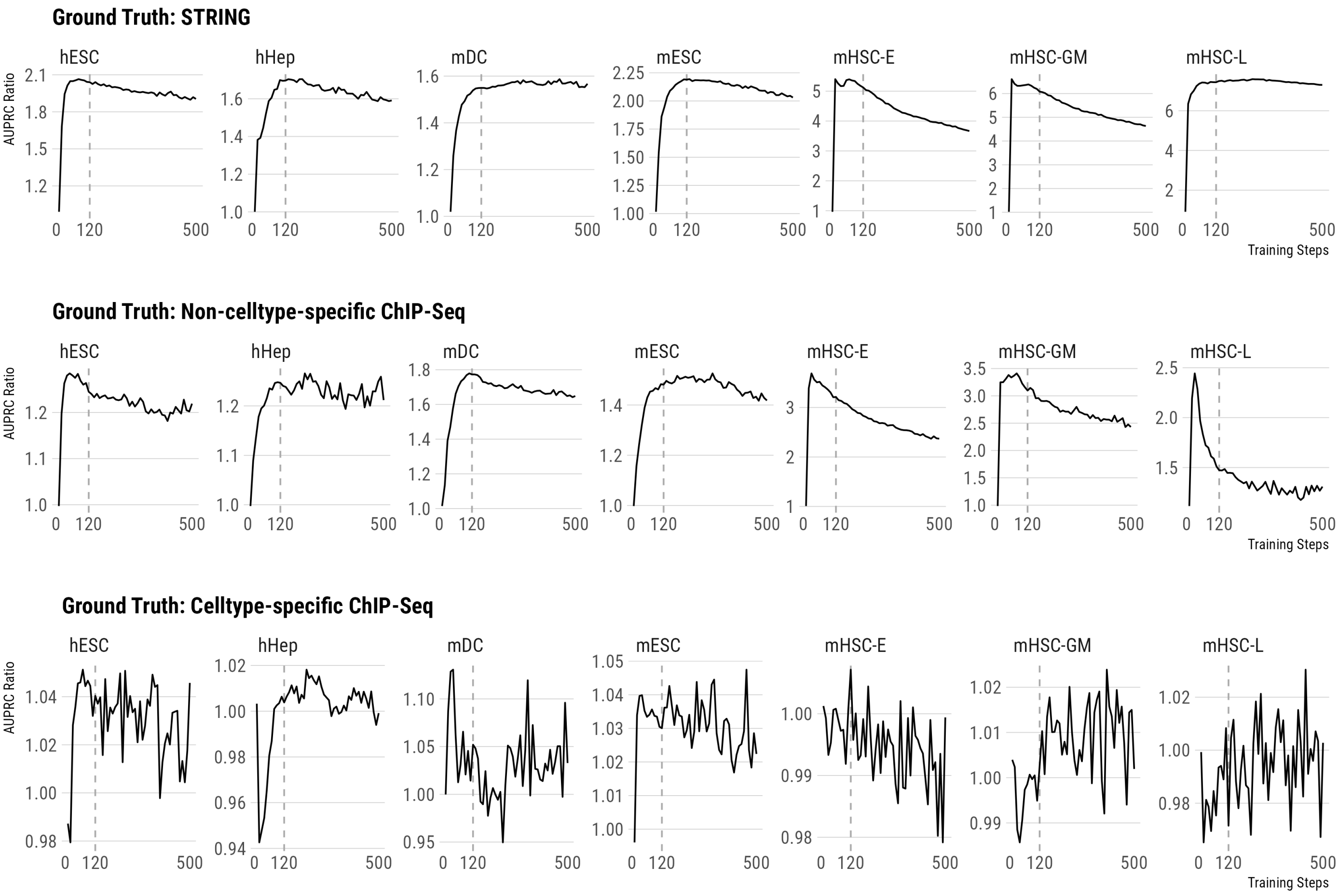


**Fig S1.** Quality of inferred GRN from DeepSEM-1x: AUPRC Ratio as a function of the number of training iterations. Quality may quickly downgrade after the performance peak. Dashed line is the recommended stopping point from DeepSEM. Note that for the celltype-specific data sets, performance is particularly volatile.

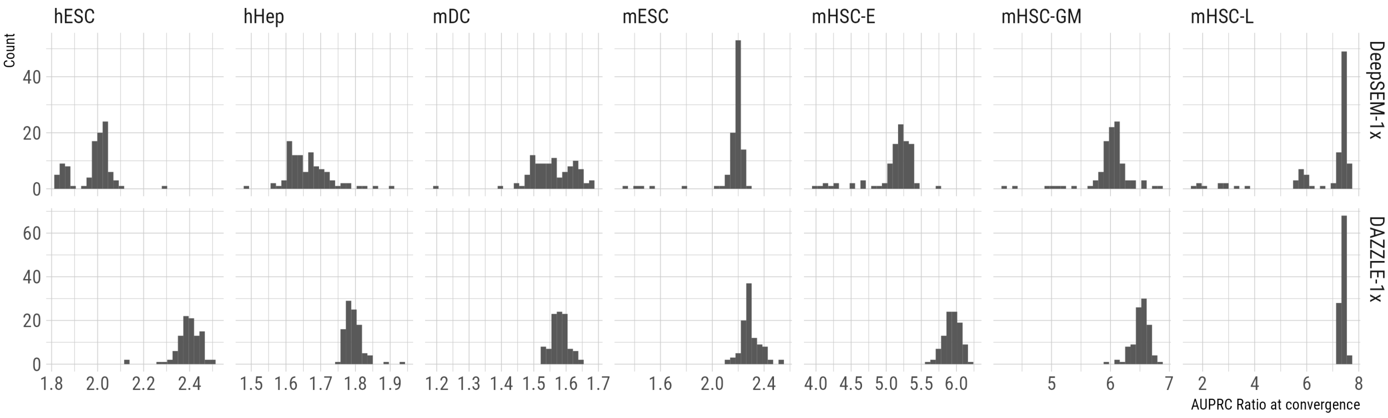


**Fig S2.** Distribution of 100 runs of DAZZLE-1x and DeepSEM-1x on BEELINE evaluated using the STRING network. Results from DAZZLE-1x tend to be more stable than results from DeepSEM-1x.
